# Trial-by-trial predictions of subjective time from human brain activity

**DOI:** 10.1101/2020.01.09.900423

**Authors:** Maxine T. Sherman, Zafeirios Fountas, Anil K. Seth, Warrick Roseboom

## Abstract

Human experience of time exhibits systematic, context-dependent deviations from veridical clock time; for example, time is experienced differently at work than on holiday. Here we test the proposal that differences from clock time in subjective experience of time arise because time estimates are constructed by accumulating the same quantity that guides perception: salient events. Healthy human participants watched naturalistic, silent videos of up to ∼1 minute in duration and estimated their duration while fMRI was acquired. We were able to reconstruct trial-by-trial biases in participants’ duration reports, which reflect subjective experience of time (rather than veridical clock time), purely from salient events in their visual cortex BOLD activity. This was not the case for control regions in auditory and somatosensory cortex, despite being able to predict clock time from all three brain areas. Our results reveal that the information arising during sensory processing of our dynamic environment provides a sufficient basis for reconstructing human subjective time estimates.

## Introduction

How do we perceive time in the scale of seconds? We know that experience of time is characterized by distortions from veridical “clock time” (1). These distortions are reflected in common expressions like “*time flies when you’re having fun*” or “*a watched pot never boils*”. That our experience of time varies so strongly in different situations illustrates that duration perception is influenced by the content of sensory experiences. This is true for low level stimulus properties, such as motion speed or rate of change (2–4), mid-level properties like complexity of task (5), and more complex natural scene properties such as scene type (e.g. walking around a busy city, the green countryside, or sitting in a quiet office; (6, 7)). It is also well-established that perception of time differs if attending to time or not (7, 8). That disruptions in time experience (i) arise across these different levels of stimulus complexity and (ii) are based on internal properties of the perceiver (such as what they are attending to) suggests that an approach that considers what is common across the hierarchy of perceptual processing, not just at a single level, is required. Further, while many studies have attempted to find a one-to-one mapping between single, simple stimulus features and time perception (e.g. speed or temporal frequency (2–4)), natural scenes contain varying proportions of any single feature and these proportions will vary over time. Therefore, to model subjective time perception on the scale of natural stimulation will clearly require an approach that jointly considers the contributions of these different features.

We recently proposed (7,9,10) that the common currency of time perception across processing hierarchies is change. In principle, this is not an entirely new idea, with similar notions having been suggested in philosophy (11) and in the roots of cognitive psychology of time (5,12,13). However, in this more recent proposal, there is a strong distinction in that change is not considered only as a function of changes in the physical nature of the stimulus being presented to the observer, but rather change is considered in terms of how the perceptual processing hierarchy of the observer responds to the stimulation.

Roseboom and colleagues (9) used a deep convolutional neural network that had been trained to classify different images as proxy for human visual processing. This study reported that simply by accumulating salient changes detected in network activity across network layers it was possible to replicate biases in human reports of duration for the same naturalistic videos. This finding supported the proposal that activity in human perceptual classification networks in response to natural stimulation could provide a sufficient basis for human time perception.

While the neural classification network used in the previous study provided a reasonable stand in for human visual processing, demonstrating at least some of the useful functional properties of human visual processing hierarchy (14, 15), fully assessing the above proposal naturally requires neural as well as behavioral evidence from human participants. Here, we put this proposal to a considerably stronger test, using a pre-registered, model-based analysis of human functional neuroimaging (BOLD), collected while participants estimated duration of silent videos. In support of our proposal, we found that the model-based analysis could produce trial-by-trial predictions of participants’ subjective duration estimates based only on the dynamics of their multi-layer visual cortex BOLD while they watched silent videos. Control models applied to auditory or somatosensory cortex could produce reasonable estimates of veridical time (i.e. clock time), but these models did not predict participant subjective reports. Our model is, to our knowledge, the first that can predict trial-by-trial subjective duration reports purely from measured human brain activity during ongoing naturalistic stimulation.

## Results

Using functional magnetic resonance imaging (fMRI) and a pre-registered preprocessing and model-based analysis pipeline (osf.io/ce9tp), we measured BOLD activation while 40 human participants watched silent videos of natural scenes (8-24 seconds each) and made duration judgements on a visual analogue scale (see Fig. 1A). Half of the videos depicted busy city scenes with many salient events, and the other half, office scenes with very few. We reasoned that if subjective time is constructed from the accumulation of salient events in sensory cortex, then videos containing more salient events (city scenes) should be judged as lasting longer relative to videos with fewer (office scenes). All inferential tests reported were preregistered unless specified otherwise.

**Figure 1.**
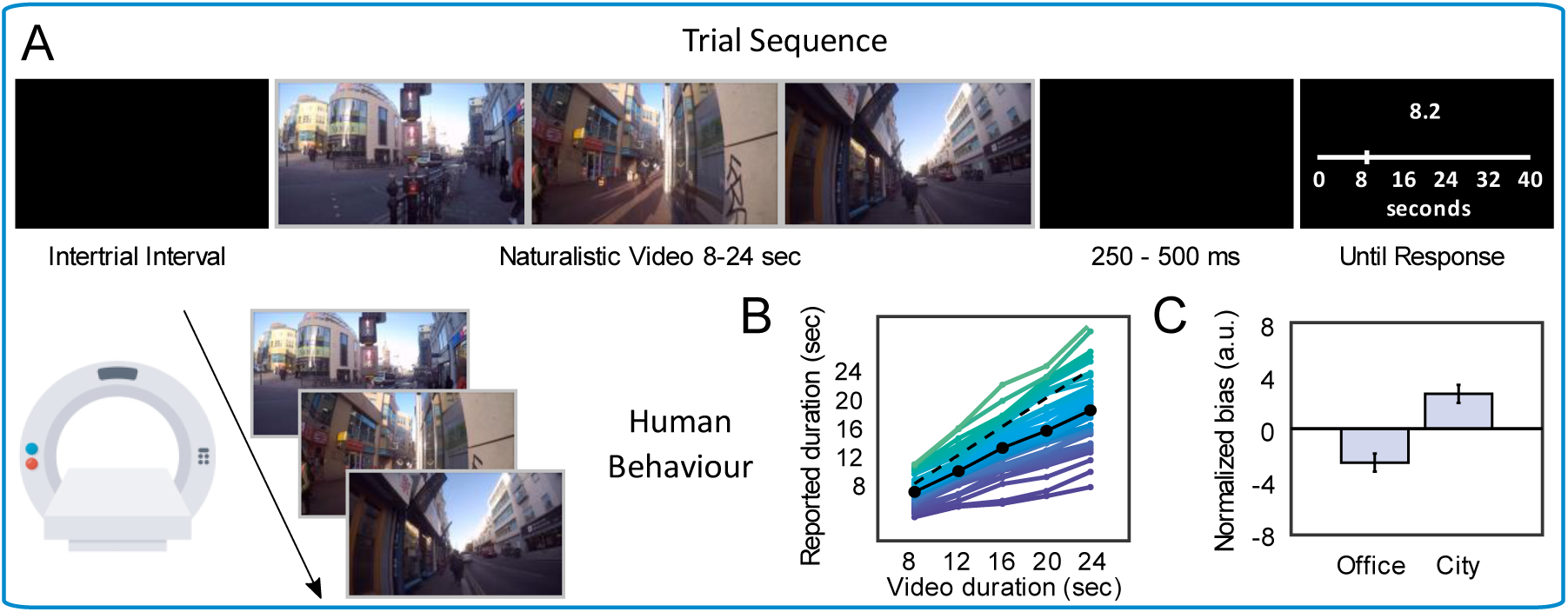
Trial sequence and human behavioral results. **(A)** Participants viewed naturalistic videos (8- 24 seconds in duration) of walking around a busy city or sitting in a quiet office while in the MRI scanner and reported the duration using a visual analogue scale. **(B)** Participant-wise relationship between report and duration (colored lines), mean relationship (solid black line), and the line of unity (dashed line). **(C)** Relative under-/over-estimation of duration by human participants for office/city videos. Error bars represent +/- within-subject SEM.

Participants’ bias towards under- or over-reporting of duration was quantified using a (pre-registered) normalized bias measure. For each of the *k* trials in which the veridical duration was *t*, the bias on trial *k* was the report on trial *k*, minus the mean report for that duration category, divided by the mean report for that category:

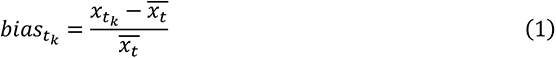

Positive/negative values mean that individual duration reports were over-/under-estimated relative to the participant’s mean for a given veridical video duration. Therefore, normalized bias here reflects participant specific response patterns that are independent of clock time.

### Behavioral reports are biased by scene type

Participants could estimate duration well, as indicated by a strong correlation between presented (veridical) and reported (subjective) durations for each subject both when computed trial-by-trial (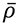 = 0.79 ± 0.10), and when averaged within duration categories (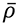 = 0.96, Fig. 1B). As predicted, durations of city scenes were relatively over-estimated and office scenes under-estimated, M_diff_ = 5.18 ± 1.36, 95% CI [1.81, 8.65], *t_39_* = 3.09, *p* = 0.004, d = 0.50, BF_H(0,10.5)_ = 33.8, confirming that natural scenes containing a higher density of salient events do indeed feel longer (Fig. 1C). Note that this result shows that the *amount* of experienced time was lower for office videos, not necessarily that time *passed faster* for office videos.

### Estimates generated by an artificial network model are biased by scene type

It has previously been shown that estimates of duration based on changes in the activation across the hierarchy of an artificial image classification network can replicate human-like biases in duration reports for naturalistic stimuli (7, 9). Here we tested whether the effect of scene type for the stimuli used in our experiment and shown by our participants (Fig. 1C) could be reproduced by this artificial perceptual classification network approach.

As in the previous study (9), we fed the same video clips that participants had viewed to a pre-trained (i.e. not trained on our stimulus set) hierarchical image classification network, AlexNet (16). For each network node, we computed frame-to-frame Euclidean distances in network activity. Then, separately for each network layer, each distance – or change in activation - was categorized as salient or not. Note that a salient event is not necessarily *psychologically* salient; it is simply a relatively extreme change in dynamics. This was achieved using an attention threshold with exponential decay that simply determined whether the change in node activation (the Euclidean distance) was sufficiently large to be deemed salient (see Methods). Following models of episodic memory (17) these moments are called ‘salient events’. Salient events were accumulated at each layer and converted to estimates of duration in seconds via multiple linear regression, by mapping the number of accumulated salient events to the *veridical*, not *reported* durations (Fig 2A). This means that the model is attempting to reproduce clock time duration based on the input, rather than the more trivial task of training the model to directly reproduce human estimates. Therefore, any human-like biases in estimates can be attributed to the behavior of the network in response to the input stimuli, and not simply to the model being trained to specifically reproduce human biases.

**Figure 2.**
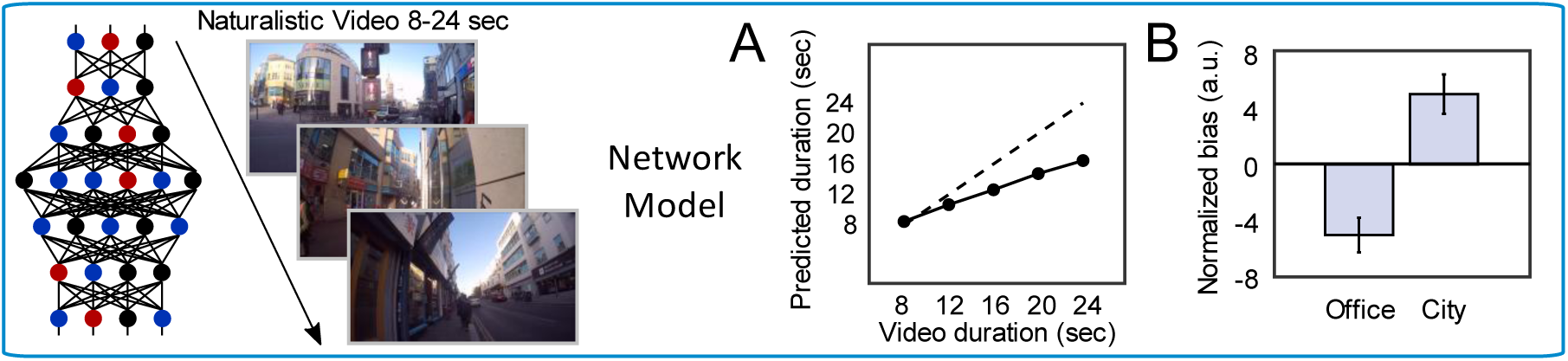
Artificial network model results. The same naturalistic videos (8-24 seconds in duration) that human participants viewed were input to our image classification network-based model to generate estimates of duration. **(A)** Relationship between veridical and model-predicted durations for this model, trained on accumulated salient events in video frames (solid line). The dashed line is the line of unity. **(B)** Relative under-/over- estimation of duration for office/city scenes for this model. Error bars represent SEM.

As was the case with human behavior, and as expected, the artificial classification network-based model produced duration reports that were significantly correlated with the video duration *ρ*(2329) = 0.73, *p* < 0.001 (Fig. 2A), indicating that the dynamics of the classifier were strongly related to clock time. More importantly, the model reproduced the pattern of subjective biases seen in human participants, despite being trained on veridical video duration. Specifically, model-produced estimates differed as a function of video type: estimation bias was greater (i.e. reports relatively over-estimated) for busy city scenes than for office scenes, M_office_ = -5.00 ± 0.66, M_city_ = 4.99 ± 0.55, 95%CI = [8.31, 11.67], *t*_2329_ = 11.65, *p* < 0.001, d = 0.48 (Fig 2B). These results demonstrate that simply tracking the dynamics of a network trained for perceptual classification while it is exposed to natural scenes can produce human-like estimates, and distortions, of duration.

### Reconstructing human-like duration reports from visual cortex BOLD

Here we put our proposal to the key test. Our proposal is that tracking changes in perceptual classification in the human perceptual processing hierarchy is sufficient to predict human trial-by-trial reports of subjective duration. Perceptual classification of visual scenes is achieved primarily in visual cortex, so to test our proposal we asked whether we could reproduce participants’ estimation biases from salient changes in visual cortex BOLD. In other words, instead of accumulating salient changes in visual *stimulation*, we accumulated salient changes in BOLD responses to that stimulation.

Coarse-level differences in BOLD were seen for both office versus city videos and for duration reports that were more biased (GLM results, see Fig. S1 and Table S4). However, this does not tell us about the relationship between duration biases and salient events in BOLD dynamics. If we can predict trial-by-trial subjective duration only from participants’ BOLD responses in visual cortex (and not in other control regions), then we will have shown that human subjective duration judgements (when viewing natural visual scenes) can be constructed from brain activity associated with perceptual processing.

To do this, we defined a three-layer visual hierarchy *a priori* predicted to be involved in processing of the silent videos (see Fig. 3 and supplementary methods). We selected regions such that lower layers reflect the detection and classification of low-level features (e.g. edge detection in primary visual cortex; V1), and higher layers, object-related processing (e.g. lateral occipital cortex; LOC). For control analyses, analogous hierarchies were built for auditory cortex and somatosensory cortex (see Table S1). Because the stimuli we used were silent videos, we predicted that only the model trained on the visual cortex hierarchy should reconstruct subjective human duration reports from accumulated salient events (see pre-registration at osf.io/ce9tp).

**Figure 3.**
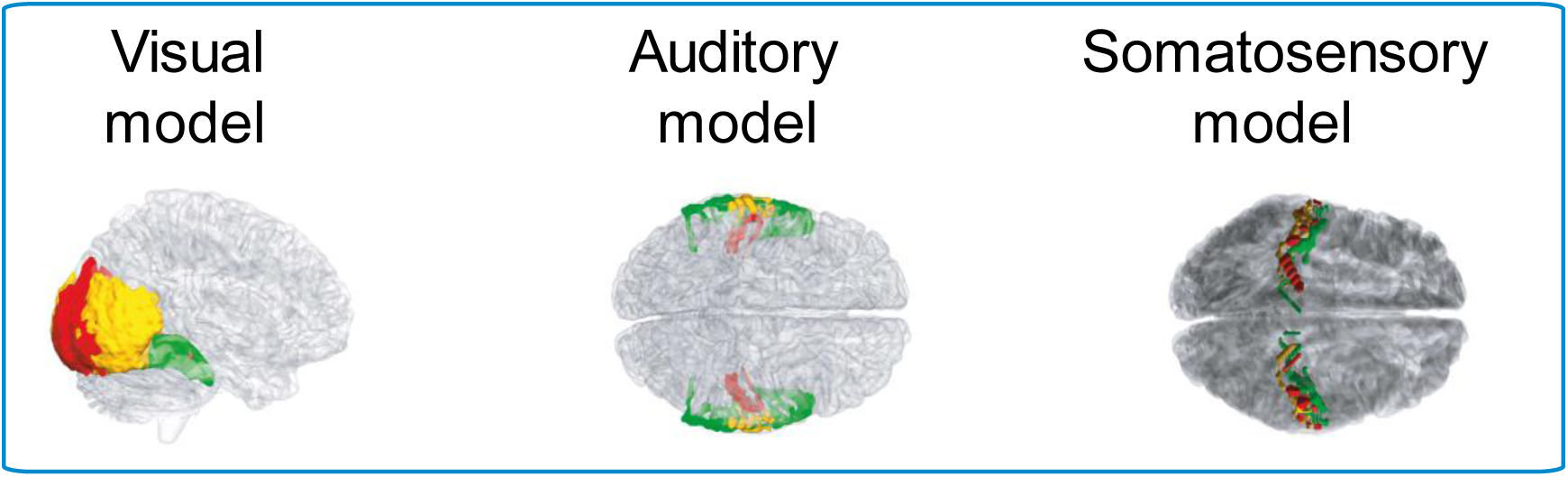
Perceptual hierarchies used for fMRI-based model analysis. Three three-layer perceptual hierarchies were defined: a visual hierarchy, an auditory hierarchy and a somatosensory hierarchy. The visual hierarchy constitutes our model-of-interest, while the auditory and somatosensory hierarchies constitute control models. The regions chosen for layers 1, 2 and 3 are colored in red, yellow and green respectively. Precise details of the regions are specified in Table S1.

We ran our key analysis in two ways: one was confirmatory (i.e. pre-registered) and one was exploratory (i.e. not pre-registered). The analysis pipeline is illustrated in Fig. 4. In both analyses, for each participant voxel-wise patterns of BOLD were extracted from each TR (slice, or time point) in each hierarchical layer. Voxel-wise changes between each TR were calculated and then summed over all voxels in the layer, resulting in one value per TR. These ‘change’ values were standardized within-participant and compared to a criterion with exponential decay to classify the change value as a salient event or not, giving us the number of salient events detected by each layer for each video.

**Figure 4.**
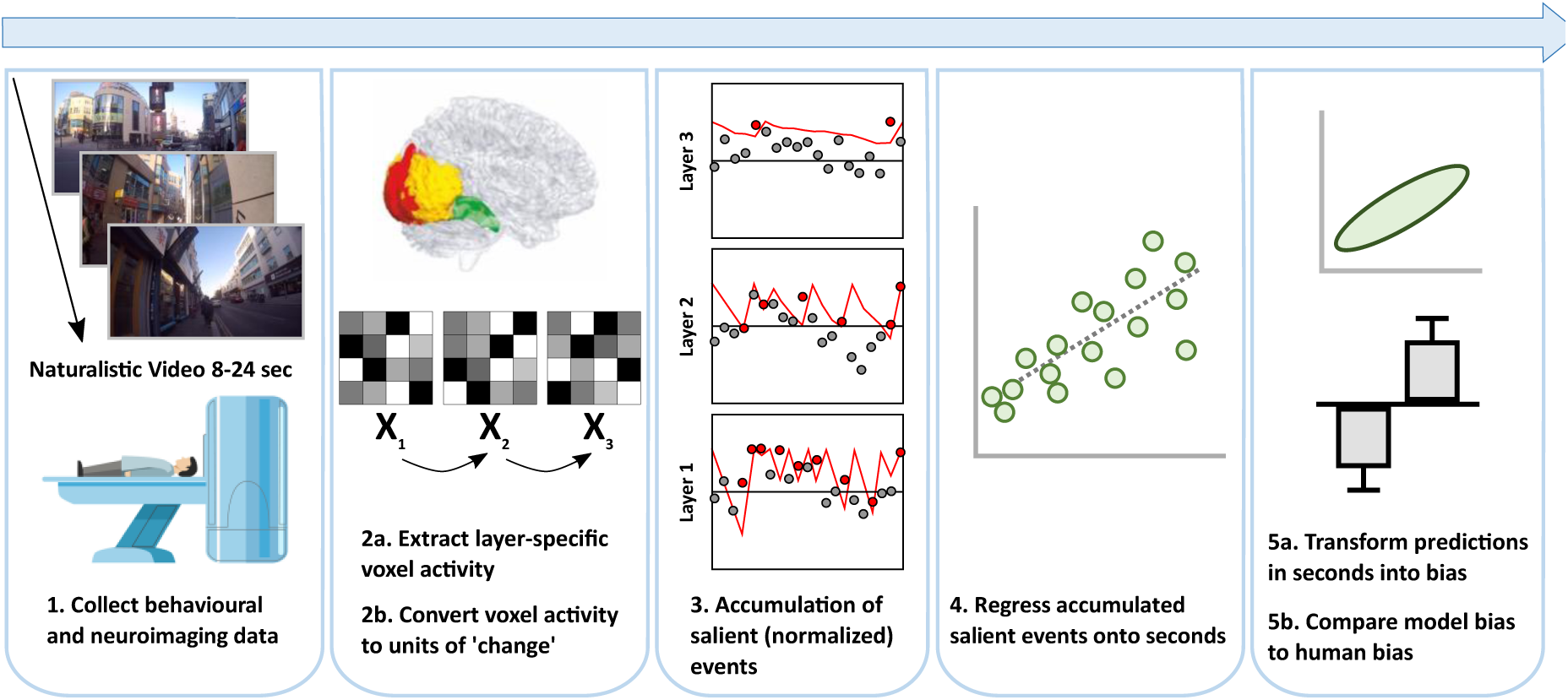
Schematic of modelling analysis pipeline. (1) Following data collection, (2) voxel-wise data was extracted and TR-by-TR changes (Euclidean distance or signed difference) computed. The example given here is for the visual hierarchy, but the same process was conducted on the auditory and somatosensory hierarchies also (see Fig. 3 and Table S1 for different hierarchies). (3) Total change in the ROI over TR was compared to an attention threshold (red line) that categorized events as salient (red dots) or not (grey dots). (4) An event was classified as salient if it took an equal or higher value than the threshold. (5) Accumulated salient events were regressed onto seconds and compared across condition and with human behavior.

For the pre-registered analysis, change was quantified as Euclidean distance (as for the artificial network model), i.e.

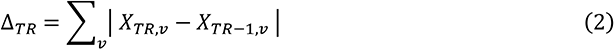

where *X_TR,v_* is activation in voxel *v* at slice *TR*. Note that (2) is mathematically equivalent to the L1 norm of the difference between BOLD at two successive TRs. However, we refer to (2) as Euclidean distances, summed over all voxels because we are proposing that the key computation here is |*X_TR,v_* – *X_TR–_*_1*,v*_| and not *X_TR,v_* – *X_TR–_*_1*,v*_.

For the exploratory analysis, we tested an alternative algorithm for quantifying ‘change’:

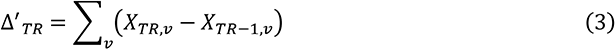

which we refer to as the signed difference. We did this because, at least in sensory cortices, BOLD may already reflect perceptual changes (18), potentially in the form of “prediction errors”. Therefore, while the model using Euclidean distance (eq. 2) as the metric of change assumes that BOLD relates *directly* to neural activity (conceptually the same as “activation” of nodes in the artificial classification network), signed difference (eq. 3) is more closely aligned with the idea that BOLD (in early sensory networks in this case) indicates (computational) *prediction error*. Euclidean distance can only be positive valued (0 or above), while the signed difference can be positive or negative in value (above or below 0).

We then used support vector regression with 10-fold cross-validation to predict veridical (i.e., not subjective/reported) video durations from accumulated salient events in layers 1, 2 and 3. This converted the accumulated salient events in each layer to model duration “reports” in seconds so that they could be compared with human reports that were also made in seconds. This step was only necessary for comparing model and human reports – as can be seen in Fig. 5, accumulated salient events (defined according to Eq. 3) in visual cortex distinguished between video type prior to their transformation into seconds.

**Figure 5.**
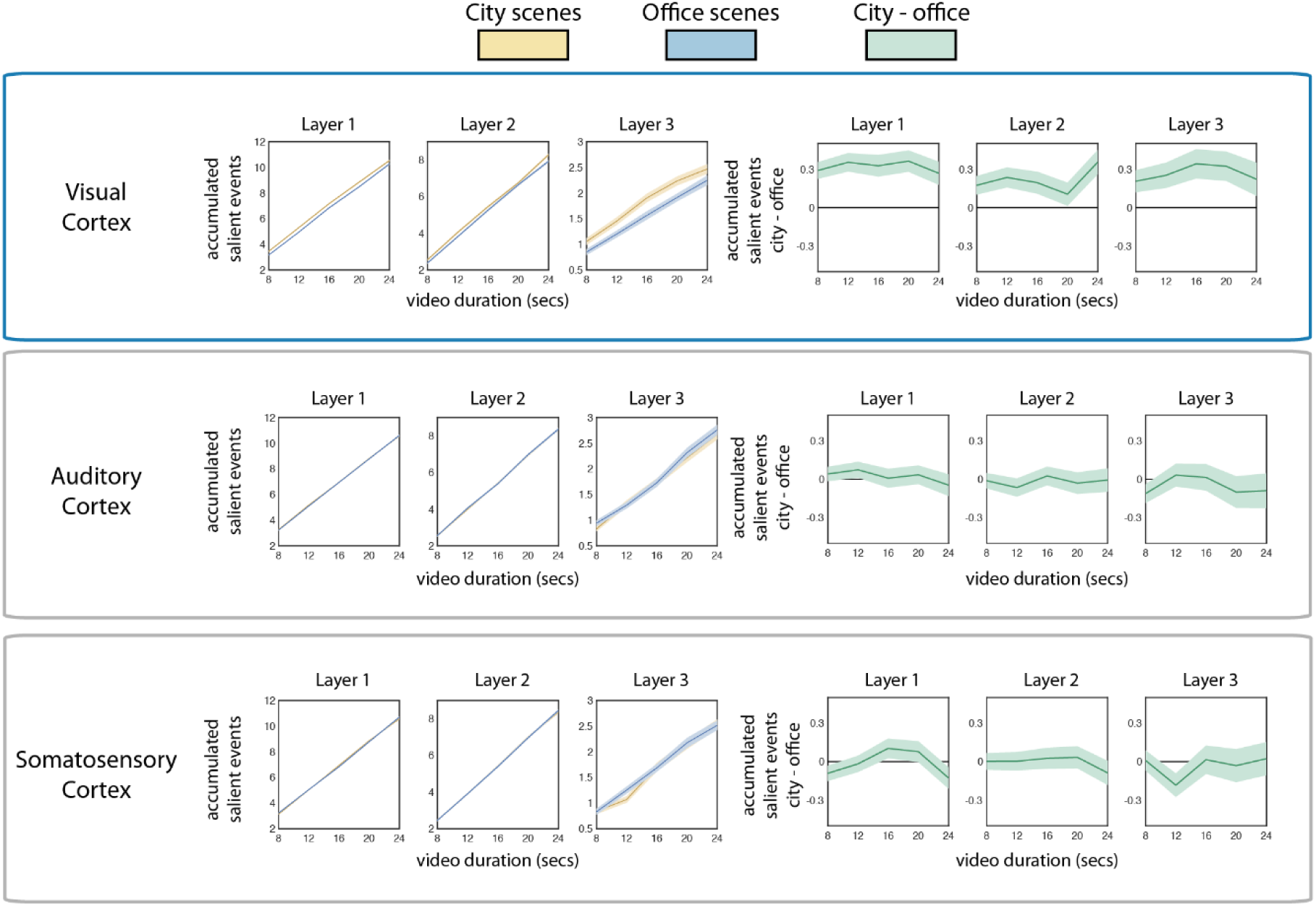
Accumulated salient events over video types, brain areas (rows) and layers (columns). The three leftmost columns plot the mean (+/-SEM) number of accumulated salient events in each layer and brain region as a function of city (blue lines) or office (yellow lines) scene. Only salient events in visual cortex distinguish between office and city scenes and this holds for all three layers. For clarity, the three rightmost columns (green lines) instead plot the difference lines, with shaded bounds depicting 95% CIs. These show that only accumulated salient events in visual cortex distinguish between scenes, because only these lines are above the zero line. This means that accumulated salient events, even prior to regression into standard units, distinguish between scene type in visual cortex, but not in auditory cortex or in somatosensory cortex.

Finally, biases in model predictions were compared to participants’ estimation biases. For our pre-registered analysis, we pooled human participants’ data together to create a ‘super-subject’, by standardizing behavioral duration reports within-participant and re-computing estimation bias on the combined behavioral dataset. For the exploratory analysis, human estimation bias was computed separately for each of the 40 participants because pooling participants’ data reduced the effect of video type on (human) estimation bias (see Fig. S2B). In both cases, model predictions were generated from the pooled accumulated changes. Participant data was standardized and pooled because the use of long stimulus presentation intervals (up to 24 seconds) meant that for each participant we could only obtain relatively few trials - insufficient to complete the analyses on a purely participant-by-participant basis.

### Using Euclidean distance, estimation bias but not effects of scene type can be reconstructed from visual cortex BOLD

Following pooling into a super-subject, the z-scored reports remained correlated with video durations (Fig.S2A) but did not significantly discriminate between office and city scenes (Fig. S2B). The presented video duration could be predicted from accumulated salient events in all three models (visual, auditory, and somatosensory) to a similar degree (10-fold cross validation, 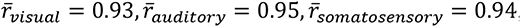, Fig. S2C-E). These results show that all models could reproduce veridical clock time – the physical duration of the presented video. The reproduction of veridical duration is trivial because, all else being equal, longer intervals will have more salient events. For this reason, our approach necessitates comparing our target model (visual cortex) with control models in other modalities (auditory or somatosensory cortex). Contrasting against control models in other modalities allows us to demonstrate that it is not simply trivial accumulation of any cortical changes over time that predicts duration, but rather accumulation of specific changes in cortical activity related to the presented content that can predict human subjective duration judgements. However, it is worth noting here that all of our models – visual cortex and the control models based on auditory or somatosensory cortex - do in fact provide reasonable estimates of veridical time. This is why our key analyses focus on reproducing the subjective *biases* present in the reports of human participants, since it is these biases that separate veridical duration from subjective duration.

Our key, pre-registered hypothesis was that only the visual cortex model would be able to reproduce participants’ duration *biases*. Supporting this, only the model trained on visual salient events significantly reproduced the super-subject’s estimation biases trial-by-trial, β_2328_ = 1.51, *p* = 0.015; the models trained on salient events in auditory cortex, β_2328_ = 0.87, *p* = 0.141, and somatosensory cortex, β_2328_ = 0.30, *p* = 0.339, did not (Fig. S3). Not only was the visual cortex regression coefficient a significant *predictor* of behavioral report, the visual cortex regression model was also a *better fit* to the trial-by-trial behavioral biases than the auditory or somatosensory cortex models (Fig. S4). These results mean that biases in subjective estimates of time can be predicted from neural activity associated with modality-specific perceptual processing. The processing is modality-specific because the video stimuli were silent, with no auditory or tactile stimulation.

While the visual model could reproduce participants’ trial-by-trial biases, it did not reproduce the effect of video type (overestimation of duration for city scenes) despite a numerical trend in the predicted direction, M_diff_ = 0.19 ±13.96, 95%CI = [-0.94, 1.33], *t_2329_* = 0.33, *p* = 0.739, d = 0.01 (Fig. S2F). The control models did not reproduce the effect of video type either (auditory: M_diff_ = -0.33 ±12.29, 95%CI = [-1.32, 0.67], *t_2329_* = -0.64, *p* = 0.522, d = -0.03, somatosensory: M_diff_ = 0.16 ±13.09, 95%CI = [-1.23, 0.90], *t_2329_* = -0.30, *p* = 0.762, d = -0.01, see Fig. S2G-H). Note that these *t-*tests were not pre-registered.

### Using Signed Difference, estimation bias and effects of scene type can be reconstructed from visual cortex BOLD

Next, we analyzed the biases predicted from the exploratory model, in which salient events were determined from signed differences in voxel activity. Again, presented video duration could be (trivially) predicted from salient events in all three exploratory models to a similar degree (10-fold cross validation, 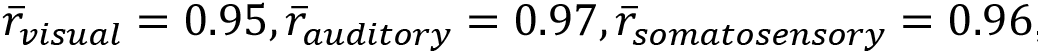, Fig 6A-C). However, using this definition of salient event, linear mixed models revealed the visual model biases *did* strongly discriminate between office and city scenes, M_diff_ = 3.75 +/- 0.23, χ^2^(1) = 85.06, *p <* .001 (Fig. 6D).

**Figure 6.**
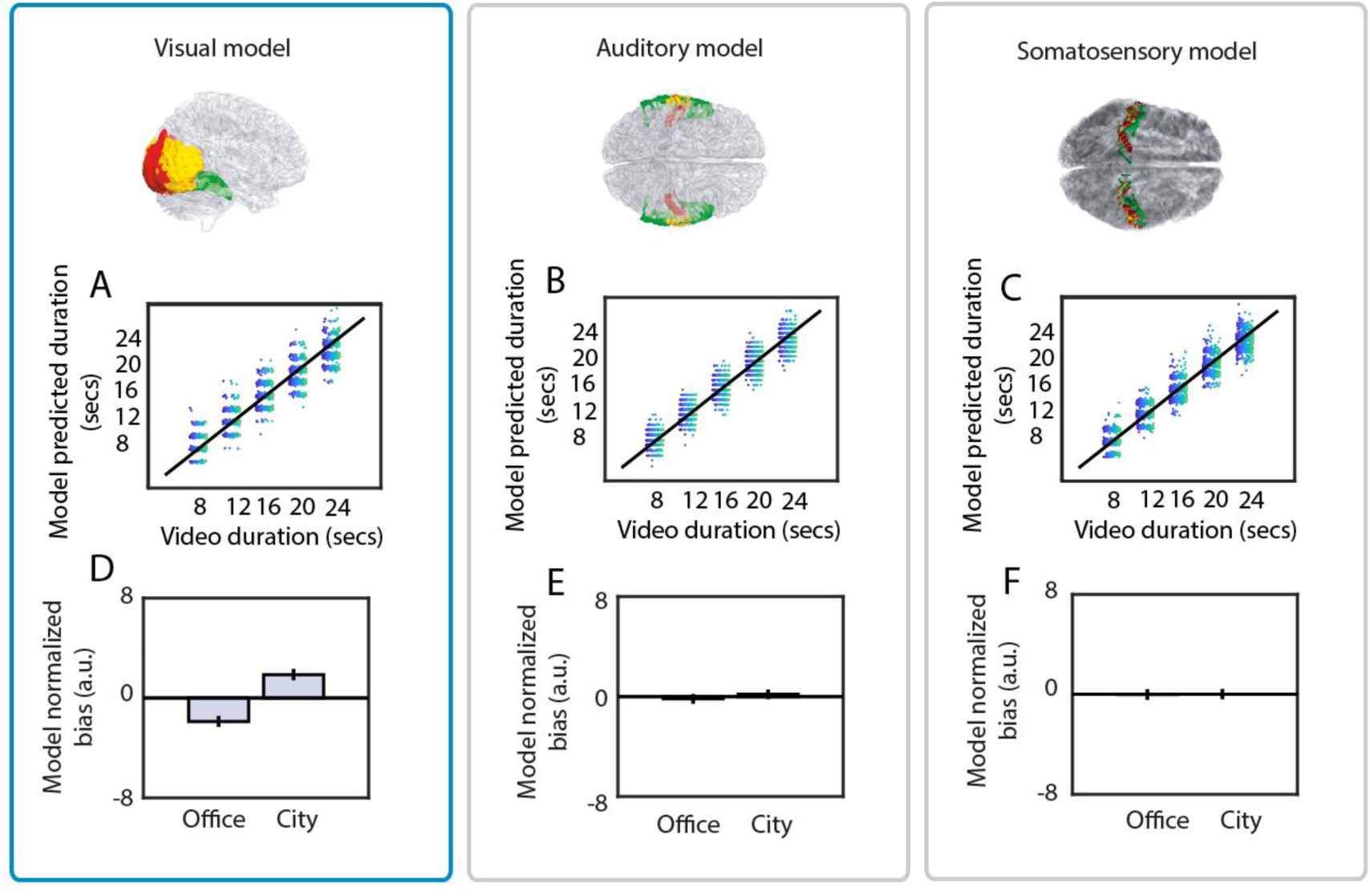
Computational neuroimaging analysis. **(A-C)** Trial-by-trial association between veridical video duration and model-predicted duration reports obtained from the visual, auditory and somatosensory models. Different dot colors represent different participants, and each dot is data from one trial. **(D-F)** Mean model-estimated normalized bias as a function of video type for the visual, auditory and somatosensory models. Error bars represent +/- within-subject SEM.

Visual model biases also remained correlated with participants’ trial-by-trial biases, β = 0.02 ±0.008, χ^2^(1) = 5.62, *p* = 0.018. This association is visualized in Fig. 7A by plotting mean model bias as a function of 25 quantiles of human normalized bias. The association held under a wide range of reasonable attention threshold parameters (Fig. 7B), meaning that model performance in reproducing participant duration reports was robust to how salient events were categorized. Again, the visual model out-performed control models in predicting normalized bias (Fig. S5). While the model trained on accumulated visual cortex salient events reproduced human behavior, biases from exploratory models trained on auditory and somatosensory salient events did not: they neither discriminated video type (M_diff_ = 0.36 ±0.19, χ^2^(1) = 0.43, *p* = 0.514, M_diff_ = 0.02 ±0.21, χ^2^(1) = 0.46, *p* = 0.499 respectively, see Fig 6E,F), nor predicted trial-wise human normalized bias (β = -0.003 ± 0.006, χ^2^(1) = 0.20, *p* = 0.652, β = 0.002±0.007, χ^2^(1) = 0.11, *p* = 0.740 respectively, Fig. 7C,E). In none of the auditory or somatosensory layers were there more salient events when watching city than office videos (Fig. 5, middle and bottom rows). Taken together, these results underline the specificity of visual cortex activity in predicting subjective time for silent videos.

**Figure 7.**
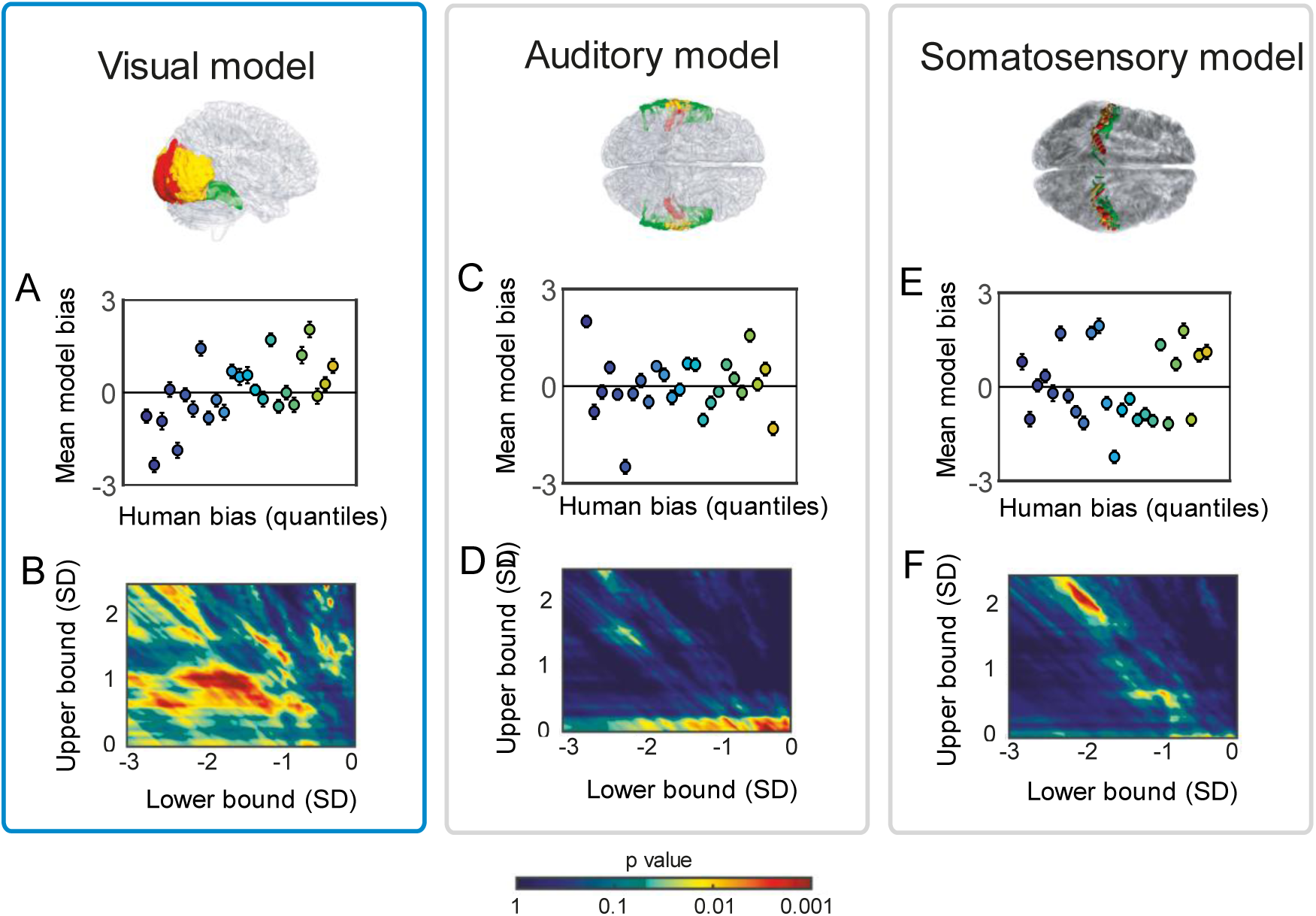
Predicting trial-by-trial subjective time from human BOLD. (**A)** Mean normalized bias for the model trained on visual cortex activity, as a function of 25 quantiles of human bias. Colors represent x-axis values. Results show a positive association between human biases and model- predicted biases. **(B)** Heat map depicting p-values for the association between human bias and (visual cortex) model bias, as a function of minimum (x-axis) and maximum (y-axis) criterion values. Dark colors represent regions where the association was non-significant at α_0.05_ or negative. Consistent results are found under a wide range of reasonable parameter values. **(C,E)** As for panel A, but for auditory and somatosensory cortex respectively. There is no association between human bias and model biases. **(D,F)** As for panel B, but for auditory and somatosensory cortex respectively. No significant positive correlation is found under alternative reasonable parameter values.

### Subjective time perception is based on perceptual processing of stimulation, not stimulus changes directly

It might be suggested that because we used naturalistic video stimuli rather than simplified, abstract stimuli, our approach is confounded by basic stimulus properties – perhaps only the low-level stimulus change present in the stimuli is driving participant and model estimation. This concern is both conceptually implausible and empirically unsubstantiated. Attempting to model human time perception based on only one type of stimulus change at a single level is unlikely to work for the reasons mentioned in the Introduction, namely that changes at higher or lower-levels of the processing hierarchy can be seen to contribute to distortions in time perception. Empirically, we examined pairs of trials where the presented scene differed (city vs office) and where the physical difference in pixel-wise changes in the videos differed greatly (up to 231 times more physical change in a city than office scene), despite reported human durations being almost identical (within 2.5% of each other). For each video clip presented to participants, we calculated the mean physical frame-by-frame change in each video as:

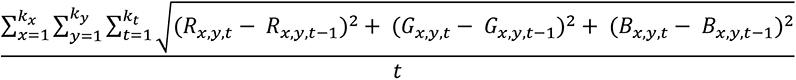

where *t* is a frame, each pixel is an (R,G,B) triplet of intensity values, and there are k_x_ x k_y_ pixels. This equation simply states that we calculated frame-to-frame Euclidean distance for each pixel separately, then summed over all pixels and timepoints. For interpretability, we divided by the number of frames so that longer videos didn’t necessarily take higher values. To identify trials where experienced durations were similar despite differing physical stimulation, we took all pairs of trials where the following conditions were met:

1. Both trials came from the same participant
2. The pair consisted of one office and one city scene
3. The log ratio between the reported durations for the two videos did not exceed 0.025, i.e. |log(report_1_/report_2_)| <= 0.025 was satisfied.

As can be seen in Fig. 8A, there were many instances in our data where the basic stimulus properties changed at a vastly higher rate in one video than another (100s of times more change in some city videos versus office videos) yet participants reported these videos as being of a very similar length (note the very narrow range of y-axis values). Despite there being clear overlap in duration reports for many instances of the different scene types, there was clearly an overall difference in duration estimation by scene (see Fig. 1C). We repeated the above process exactly but seeking trials that differed in the visual cortex model predictions instead. The results are plotted in Fig. 8B, and again, show that two trials with substantially different stimulus properties can nonetheless be associated with similar duration estimates predicted from visual cortex BOLD.

**Figure 8.**
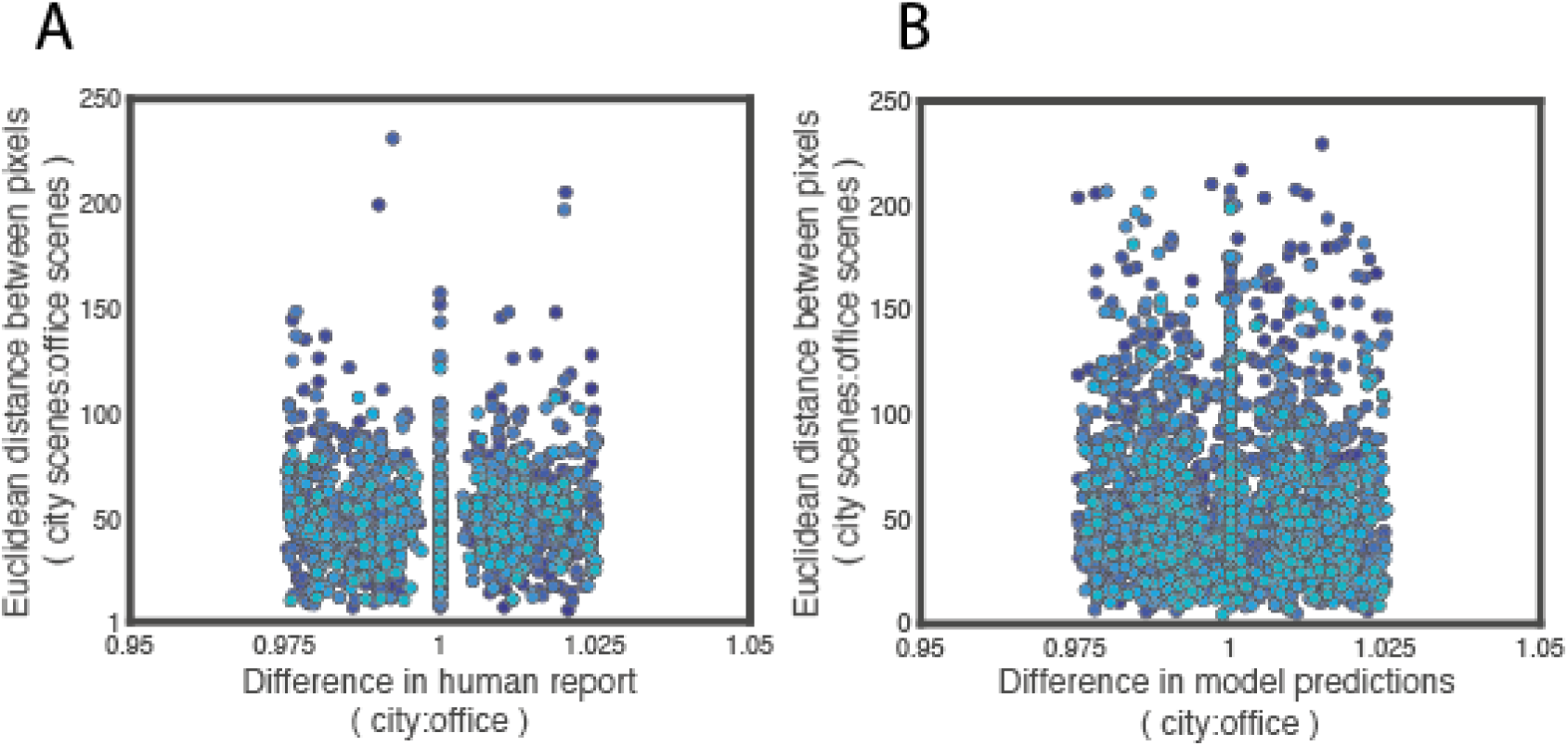
Dissociation between human and model duration estimates and stimulus properties. **(A)** Each dot is the pixel-wise difference for a pair of city and office trials presented to a given participant where the log ratio between the reported durations for the two videos did not exceed 0.025 (i.e. they were estimated as very similar). Dot colors represent different participants. This figure shows that there were many trials in our data where, despite being very different from another trial (up to 100s of times different in pixel-wise distance), the two trials were estimated as almost the same duration. **(B)** As for A, but where visual cortex model predictions were similar. Again, this panel shows that there were many trials in our data where the visual information differed greatly, but the duration estimate predicted by our visual cortex model was almost the same.

These results are what one would expect given that the stimuli we used are naturalistic videos: namely, there is a wide distribution of physical changes at the most basic level because the videos contain real scenes filmed in the world, rather than contrived abstract stimuli typically used in studies of time perception. These basic physical changes are associated with different types of natural events – a bus going past on the adjacent street, a person walking in front of the camera, a person appearing in view in a largely empty office – but not through a direct one-to-one mapping.

These two results demonstrate that it is not the changes detectable in the videos themselves that is key, but the response of perceptual classification networks - artificial or biological - to that stimulation that allows us to predict human reports of subjective time. That our approach tracks these changes across a hierarchy of processing is a feature that allows the model to deal with time perception on a natural scale. It is not a confound to be eliminated by considering only video stimuli perfectly matched for their lowest-level properties (i.e, mean pixel-wise distance) – since doing so would render such stimuli uninformative about naturalistic perception.

## Discussion

We have shown that subjective estimates of duration can be constructed on a trial-by-trial basis from salient events in sensory cortex activity, where salient events are defined as relatively large changes in the neural responses to sensory stimulation across a perceptual hierarchy. Importantly, participants are not necessarily conscious of the events (because they are events in the processing dynamics rather than in the stimulus), and salient events are not necessarily *psychologically* salient. In this study, for which stimuli were silent videos, successful prediction was obtained only for models trained on salient events in visual cortex BOLD, and not for control models based on somatosensory or auditory cortex BOLD. While we could trivially reconstruct veridical clock time from activity in all three sensory regions (because those regions exhibited dynamic neural activity that is correlated with veridical duration), only the information extracted by the stimulus-relevant sensory model - the visual model - was related to subjective duration estimates. Our results were robust under a wide range of model parameter values (Fig. 7B,D,F), and, in combination with results from the perceptual classification network model and previous findings (19, 20), support the idea that human time perception is based in the neural processes associated with processing the sensory context in which time is being judged.

### A novel approach to modelling subjective time

Our model is the first which is able to predict trial-by-trial subjective reports of duration based on neural (BOLD) activity during naturalistic stimulation, and in so doing, advances our understanding of the neural basis of time perception. Our approach is conceptually related to a study by Ahrens and Sahani (21), who proposed that subjective time perception is constructed from estimates of input dynamics (akin to sensory input) and knowledge of the temporal structure of the input (second order input statistics), and presented an inference model that could account for several behavioural results. An important difference between their work and ours is that statistical “knowledge” in our model relates to knowledge of the perceptual classification network state. y contrast, knowledge in hrens and Sahani’s model relates to prior temporal structure. This means that while Ahrens and Sahani propose a model dependent on processes dedicated to tracking temporal properties, we do not. Our results demonstrate that such knowledge is not strictly necessary for generating human-like duration estimates for natural stimuli.

### Explicitly linking sensory content and subjective duration

In our approach, the neural processes that are engaged in the processing of sensory content (perceptual classification processes) are the same as those used to build estimates of time. In this way we provide an intuitive link between sensory content and subjective duration. Our conclusion is in support of the idea that time perception depends on distributed mechanisms (22), but that in each case subjective time is naturally linked to sensory content by virtue of being determined by those content-related processes. Our work provides an important step from previous studies on the topic. It is common for studies to use highly constrained stimuli such as luminance contrast discs, Gabors, or random dot fields. The high level of control over stimulation in these studies is to attempt to provide a one-to-one mapping between a single stimulus feature (e.g. contrast intensity or temporal frequency) and time perception. However, since natural scenes contain a wide variety and combination of simple features, some of which are coded for in overlapping neural populations, it is necessary to consider the joint contributions of these features. By using a metric based on salient events across the perceptual processing hierarchy, our approach accomplishes this task, allowing us to model natural stimulation in a way where tracking only a single stimulus feature would fail (e.g. tracking only luminance contrast intensity in natural scenes would not predict time perception for scenes where little luminance change occurs; tracking only motion energy would not predict time perception for scenes where there is little movement).

In the time perception literature there are often appeals to the influence of factors like attention (23); or emotion (24, 25) – sometimes used as the basis for rejecting the proposals of earlier cognitive psychologists such as Ornstein (5). Our model provides a general basis on which to test any claims about any such influences by specifying a baseline hypothesis in each case – the dynamics of the relevant sensory cortex (e.g. visual, auditory, interoceptive, etc) are sufficient to construct subjective duration estimates for that context. For example, regarding the potential influence of emotion on time perception (24, 25), the degree to which stimulus-driven differences in network activation underlie any differences in duration estimates remains to be established. However, even if influences of emotion on time perception arise from internally generated sources (rather than processing of external stimulation), this influence may still be reflected in differences in measurable activity of perceptual classification networks (via e.g. BOLD) and therefore our model would reproduce differences in human duration reports. These possibilities are testable hypotheses, made available by our modelling approach.

### Time perception for non-visual or multisensory cases

While we only tested whether subjective duration for visual stimuli could be constructed from salient events in visual cortex, we expect that salient events from auditory cortex would predict subjective time in auditory-only contexts, and likewise other modalities (26). Outside the laboratory we judge time in multisensory contexts and can estimate duration when our eyes are closed or even if clinically deaf. These observations do not expose a weakness of our approach, but generate specific and testable claims that require additional study to fully evaluate: does our model approach work equally well in other (combinations of) modalities? We see no why our proposal would fail for other cases, and the success of similar network architectures for interpreting sensory processing in the auditory domain (27) supports this position. Furthermore, because we define ‘salient events’ as events in the dynamics of the perceptual classification process rather than in the external world, non-external stimulation like visual imagery could also contribute to experience of time.

It could be suggested that our approach only reproduces stimulus-driven biases rather than providing a general basis for time estimation because without stimulus input our model would have no “activity”. This critique would be valid for the artificial network-based model in section *Estimates generated by an artificial network model are biased by scene type* but cannot be applied to the BOLD-based model because visual cortex activity remains present even when there is no sensory input (see also above point about imagery).

### Biological plausibility

Our data do not speak to the question of *how* perceptual classification is achieved by the brain, and our results do not depend on the answer. Whether the classification network used here (AlexNet) is closely matched to biological vision in how it processes information not relevant here (see (14,15,28)) because the algorithmic approach to estimating duration from network activity (in either AlexNet or estimated from BOLD in humans) produces outcomes consistent with the patterns seen in human subjective reports of time. The crucial assumption is simply the existence of a hierarchical, specialized system for perceptual classification - the common interpretation of primate ventral visual stream (29–31). Given this assumption, whatever the specific computational processes underlying perception for the human brain are, the dynamics of perceptual systems implementing those processes can be used to construct subjective duration. This conclusion is best demonstrated by the fact that our model produced estimates consistent with subjective biases in human reports regardless of whether applied to activation patterns of AlexNet or to BOLD patterns recorded from human participants.

### Predictive processing as a potential mechanistic basis for time perception

We tested two metrics that could be used by the brain to link sensory content and time on a moment-to-moment basis: Euclidean distance (pre-registered) and signed difference (exploratory). While the former assumes that BOLD activity indexes some raw quantity associated with sensory inputs, the latter assumes that BOLD already indexes change in sensory input, for example as perceptual prediction error. In our data, subjective duration was best reconstructed using signed difference: although both metrics generated duration estimates that correlated with human reports, only the signed metric differentiated video type. The superiority of signed difference in predicting subjective time is consistent with (but not evidence for) the view that BOLD already indexes detected environmental changes. This is in line with literature evidencing “surprise” or “prediction error” responses in sensory (18,32,33) and even frontal (34, 35) cortices, usually interpreted in the context of predictive processing (36) or predictive coding (27) theories of cortical function. Of course, this superiority of signed difference is not itself evidence for a role for prediction error in time perception, nor are the theories of predictive processing (36) or predictive coding (37) necessary for understanding or interpreting our results.

We also emphasize that the way in which we use “salience” and “surprise” is only tangentially, if at all, related to the psychological phenomena of something being salient or surprising. Here, salience is defined in terms of difference between successive network states (see Eqs. 1 and 4). This means our notion of salience is close to a naïve prediction error (9); naïve because the “prediction” is simply the previous network state rather than part of a prediction-update cycle (see (20)). While previous studies have suggested that predictability (38) or apparent salience (39) can affect subjective time perception (40), descriptions of “salience” and related terms at this cognitive level are not necessarily related to descriptions at the mechanistic level at which our model is articulated. Future work may wish to test whether “prediction error” defined in a mechanistic sense maps onto psychological salience or surprise, but the question is outside the scope of the present study, and is certainly not restricted to investigations of time perception.

### “Surprise”, time perception, and episodic memory

The idea that our model may be based on an index of perceptual “surprise” is nonetheless intriguing as it provides a natural link to the closely related topic of episodic memory (see (20)). In the episodic memory literature, prediction error, i.e. the difference between current sensory stimulation and expected stimulation, has been proposed as the basis for the construction of event boundaries (17,20,41) – transitions that segment some content (e.g. a cow) from some other content (e.g. a car) in continuous experience (42, 43). By emphasizing the importance of sensory content in time perception, our approach may provide a link between time perception and episodic memory that has been lost by content-free “clock” approaches. By providing a simple algorithm for how the stream of basic sensory processing is segmented into salient events, our approach may afford some insight into how low-level sensory information is transformed into the temporally sequenced form of memory associated with the activity of so-called “time cells” (44–46), linking the content of basic sensory processing with temporal properties of episodic memory within the powerful predictive coding approach (20,37,47).

## Conclusions

In summary, we provide evidence for a simple algorithmic account of duration perception, in which information sufficient for subjective time estimation can be obtained simply by tracking the dynamics of the relevant perceptual processing hierarchy. In this view, the processes underlying subjective time have their neural substrates in perceptual and memory systems, not in systems specialized for time itself. We have taken a model-based approach to describe how sensory information arriving in primary sensory areas is transformed into subjective time, and validated this approach using human neuroimaging data. Our model provides a computational basis from which we can unravel how human subjective time is generated, encompassing every step from low level sensory processing to the detection of salient perceptual events, and further on to the construction and ordering of episodic memory.

## Materials and Methods

### Participants

The study was approved by the Brighton and Sussex Medical School Research Governance and Ethics Committee (reference number ERA/MS547/17/1). Forty healthy, English speaking and right-handed participants were tested (18-43 years old, mean age = 22y 10mo, 26 females). All participants gave informed, written consent and were reimbursed £15 for their time. Sample size was determined according to funding availability.

### Procedure

The experiment was conducted in one sixty-minute session. Participants were placed in the scanner and viewed a computer visual display via a head-mounted eyetracker, placed over a 64-channel head coil. Eyetracker calibration lasted approximately five minutes and involved participants tracking a black, shrinking dot across nine locations: in the center, corners and sides of the visual display. Eyetracking data are not used in this manuscript due to technical failure.

Following calibration, we acquired six images reflecting distortions in the magnetic field (three in each of the posterior-to-anterior and anterior-to-posterior directions) and one T1-weighted structural scan.

Finally, functional echoplanar images (EPIs) were acquired while participants performed two to four blocks (time-permitting) of twenty trials, in which participants viewed silent videos of variable length and reported the duration of each video using a visual analogue scale extending from 0 to 40 seconds (see Fig. 1A). A key grip was placed in each hand, and participants moved a slider left and right using a key press with the corresponding hand. Participants were not trained on the task prior to the experimental session.

### Experimental design and trial sequence

Each experimental block consisted of 20 trials. On each trial a video of duration 8, 12, 16, 20 or 24 seconds was presented. For each participant, videos of the appropriate duration and scene category were constructed by randomly sampling continuous frames from the stimuli built for (9). These videos depicted either an office scene or a city scene. Two videos for each duration and content condition were presented per block in randomized order. For one participant and one block, only 11/20 trials were completed giving a total of 2331 trials across the entire dataset.

### MRI acquisition and pre-processing (confirmatory)

Functional T2* sensitive multi-band echoplanar images (EPIs) were acquired on a Siemens PRISMA 3T scanner (2mm slices with 2mm gaps, TR = 800ms, multiband factor = 8, TE = 37ms, Flip angle = 52°). To minimize signal dropout from parietal, motor and occipital cortices, axial slices were tilted. Full brain T1-weighted structural scans were acquired on the same scanner using the MPRAGE protocol and consisting of 176 1mm thick sagittal slices (TR = 2730ms, TE = 3.57ms, FOV = 224mm x 256mm, Flip angle = 52°). Finally, we collected reverse-phase spin echo field maps, with three volumes for each of the posterior to anterior and anterior to posterior directions (TR = 8000ms, TE = 66ms, Flip Angle = 90°). Corrections for field distortions were applied by building fieldmaps from the two phase-encoded image sets using FS ’s TOP P function. ll other image pre-processing was conducted using SPM12 (http://www.fil.ion.ucl.ac.uk/spm/software/spm12).

The first four functional volumes of each run were treated as dummy scans and discarded. A standard image pre-processing pipeline was used: anatomical and functional images were reoriented to the anterior commissure; EPIs were aligned to each other, unwarped using the fieldmaps, and co-registered to the structural scan by minimizing normalized mutual information. Note that in accordance with HCP guidelines for multiband fMRI we did not perform slice-time correction (48). After co-registration, EPIs were spatially normalized to MNI space using parameters obtained from the segmentation of T1 images into grey and white matter, then smoothed with a 4mm FWHM Gaussian smoothing kernel. Smoothed data were used for the GLM on BOLD only; unsmoothed data were used for the brain-based modelling.

### Statistical analyses

All fMRI pre-processing, participant exclusion criteria, behavioral, imaging and computational analyses were comprehensively pre-registered while data collection was ongoing (osf.io/ce9tp/) but before it was completed. This analysis plan was determined based on pilot data from four participants, and was written blind to the data included in this manuscript. Analyses that deviate from the pre-registered analysis plan are marked as “exploratory”. Pre-registered analyses are described as “confirmatory”. Data are freely available to download at https://osf.io/2zqfu.

### fMRI statistical analysis (confirmatory)

At the participant level, BOLD responses obtained from the smoothed images were time-locked to video onset. BOLD responses were modelled by convolving the canonical hemodynamic response function with a boxcar function (representing video presentation) with width equal to video duration. Videos of office and city scenes were modelled using one dummy-coded regressor each. Each was parametrically modulated by normalized bias.

Data from each run was entered separately. No band-pass filter was applied. Instead, low-frequency drifts were regressed out by entering white matter drift (averaged over the brain) as a nuisance regressor (35, 49). Nuisance regressors representing the experimental run and six head motion parameters were also included in the first level models. Because of the fast TR, models were estimated using the ‘F ST’ method implemented in SPM.

Comparisons of interest were tested by running four one-sample t-tests against zero at the participant level for each variable of interest (video scenes, office scenes, and their normalized bias parametric modulator). Next, group-level F tests were run on those one-sample contrast images to test for effects of video type and the interaction between video type and normalized bias slope. A one-sample t-test against zero at the group level tested the slope of the normalized bias-BOLD relationship. All group-level contrasts were run with peak thresholds of p < .001 (uncorrected) and corrected for multiple comparisons at the cluster level using the FWE method. Clusters were labelled using WFU PickAtlas software (50, 51).

### Model-based fMRI (confirmatory)

Our key prediction was that subjective duration estimates (for these silent videos) arise from the accumulation of salient (perceptual) events detected by the visual system, particularly within higher-level regions related to object processing. We tested this by defining a (pre-registered) three-layer hierarchy of regions to represent core features of the visual system:

Layer 1 was defined as bilateral V1, V2v and V3v, Layer 2 was defined as bilateral hV4, LO1 and LO2, and Layer 3 as bilateral VO1, VO2, PHC1 and PHC2 (clusters are depicted in Figure 1). For each layer, masks were constructed by combining voxels from each area, using the atlas presented in (52).

To determine events detected by the visual system over the course of each video, we extracted raw voxel activity for each TR in each layer from unsmoothed, normalized EPIs. Then, for each voxel v, change was defined as the Euclidean distance between BOLD activation x_v_ at volume TR and TR-1. The amount of change detected by the layer at any time point, denoted Δ_TR_, was then given by summing the Euclidean distances over all voxels such that:

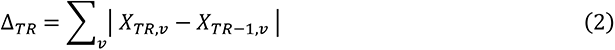

This process furnishes one value per layer for each TR of each trial for each participant. The next step was to categorize each value as a “salient” event or not and convert it to an estimate of duration using an event detection, accumulation and regression model, as presented in Roseboom et al. (9). Before converting accumulated salient changes to units of seconds, we first pooled participants’ data by z-scoring the summed events Δ_TR_ within each participant and layer. Pooling was performed to increase statistical power of subsequent regression analyses. Then, for each trial, TR-by-TR categorization of Δ_TR_ was achieved by comparing against a criterion with exponential decay, corrupted by Gaussian noise ε:

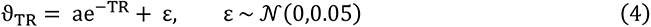

Only the parameter a took different values in each layer (see Table S2): it took larger values at higher layers. The criterion decayed with each TR until either an event was classified as salient or until the video finished, after each of which the criterion reset to its starting (i.e. maximal) point. Importantly, because the summed Euclidean distances Δ_TR_ were z-scored, the criterion has meaningful units corresponding to SDs above or below the mean. The parameter *a* corresponds to the largest z-score above which a change was classified as salient, that is, the criterion’s most conservative point. To account for potential head-motion artefacts, criterion updating ignored volumes where Δ_TR_ was greater than 2.5 (i.e. more than 2.5 SDs from the mean).

The final modelling step was to convert the BOLD-determined accumulation of salient events into raw duration judgements (in seconds). This was achieved via Epsilon-support vector regression (SVR), implemented on python 3.0 using *sklearn* (53), to regress accumulated events in each of the three layers onto the veridical video duration.

To evaluate whether the model could reproduce subjective reports of time from participants’ O activation, we converted the trial-by-trial model predictions (raw duration judgements in seconds) to normalized bias. These were then compared to a human “super-subject”: participants’ duration judgements were z-scored within participants, then all participant data were pooled and converted to normalized bias. We created a super-subject to mirror the data pooling performed before training our SVR.

Trial-by-trial normalized bias values were compared across model and human using linear regression, fitting the model:

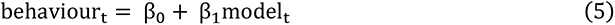

To test our *a priori* hypothesis that the model trained on visual cortex salient events positively correlates with subjective time, a (one-tailed) p-value for β_1_ was calculated via bootstrapping, shuffling the behavioural data and refitting the regression line 10,000 times.

### Control models (confirmatory)

To distinguish our proposal from the more trivial suggestion that the neural dynamics of any cortical hierarchy (or any neural ensemble) can be used to approximate elapsed clock time, simply because they are dynamic, we created two control models. While these models should all approximately reproduce clock time, the reproduced estimates should not be predictive of the specifically subjective aspects human participants’ duration estimates (i.e., their biases). Analyses for these control hierarchies followed the steps above for the primary model, though based on different sensory regions.

The first control hierarchy was auditory cortex, which has previously been implicated in time perception but whose involvement in duration judgements should not be driven by visual stimuli, as in our study. Layers 1 and 2 were defined as Brodmann Area (BA) 41 and 42 respectively, both of which are located in primary auditory cortex. Layer 3 was posterior BA22 (superior temporal gyrus/Wernicke’s rea).

The second control hierarchy was somatosensory cortex, which on our model should not be involved in duration judgements based on visual stimuli. Layer 1 was set as posterior and anterior BA 3, and layers 2 and 3 were set as BA 1 and BA 2 respectively. These Brodmann areas correspond to the primary somatosensory cortex.

Masks for these two control analyses were constructed using WFU PickAtlas atlases (50, 51). As for our empirical analyses using visual cortex, for each of the two controls we estimated the relationship between the trial-by-trial normalized bias based on the model’s predictions and based on z-scored participant data by fitting a linear regression line.

To test whether the visual cortex model out-performed the somatosensory and auditory cortex models we compared their log-likelihoods, obtained from the Matlab function *fitlm* (see Fig. S4). This evaluation of model performance was not pre-registered.

### Exploratory modelling

We also ran an exploratory (i.e. not pre-registered) set of models. This was identical to the pre-registered analysis plan, apart from the following differences:

First, we transformed voxel-wise BOLD activation X to signed (i.e. raw) rather than unsigned changes:

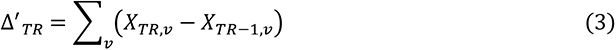

Using SVR as before, for each hierarchy we obtained model-predicted duration estimates in seconds. To avoid pooling participants’ reports together, human judgements were not standardized. Instead, for each of our 40 participants we computed human and model normalized biases from the human reports and model predictions associated with the set of videos associated with each participant. In other words, normalized bias was computed ‘within-participant’.

To test the association between video-by-video human and model bias while accounting within-participant variability we used a linear mixed model approach. Using R with the *lmer* and *car* packages, we fit the following random-intercept model:

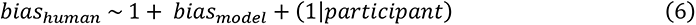

To determine whether model (*bias_model_*) and human (*bias_human_*) biases correlate, we used a chi-squared test (from the *car* function *Anova*) to compare eq. 5 to a reduced model without the fixed effect:

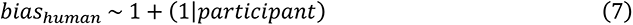

To test the effect of video type (or scene) on model normalized bias, we fit the model:

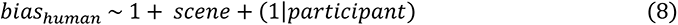

Again, we used a chi-squared test to compare Eq. 8 to the reduced model that did not include *scene* (Eq. 7)

To test whether the model trained on visual cortex events outperformed the somatosensory and auditory models, we compared the difference in AIC between the main (Eq. 6 and Eq. 8) and control (Eq. 7) models for each hierarchy (see Fig. S5).

### Robustness analysis (exploratory)

To examine the robustness of our exploratory analysis to criterion parameters we reran the above analysis pipeline under varying values of ϑ*_min_* and ϑ*_max_*. For layer 1 (where there should be most salient changes), ϑ*_min_* took 50 linearly-spaced values between 3 SD and 0 SD below the mean. ϑ*_max_* independently took 50 linearly-spaced values between 0 SD and 2.5 SD above the mean. We chose 2.5 SD because this was the highest value z-scored BOLD could take before being discarded as a head motion artefact. For each pair of ϑ*_min_* and ϑ*_max_* values for layer 1, the lower/upper bounds for layer 2 were ϑ*_min_* + 0.5 and ϑ*_max_* + 0.5 respectively. For layer 3, they were ϑ*_min_* + 1 and ϑ*_max_* + 1 respectively.

With these criteria, we obtained 250 datasets for each ROI. For each ROI and dataset, we tested the association between model-predicted bias and human bias by fitting the regression model:

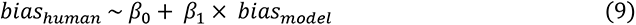

Heat maps depicted in Fig. 1 correspond to one-tailed p-values for β_1_. This robustness analysis was not pre-registered.

### Artificial classification network-based modelling

Frames from each video presented during the experiment were fed into the model presented in Roseboom et al (6). Instead of accumulating events based on changes in BOLD amplitude, salient events in the video frames themselves were detected by analyzing activity in an artificial image classification network (AlexNet)(16). We used nine network layers (input, conv1, conv2, conv3, conv4, conv5, fc6, fc7, and output, where fc corresponds to a fully connected layer and conv to the combination of a convolutional and a max pooling layer). Node-wise Euclidean distances for each node were computed, then summed over all nodes in the layer giving one value per video frame and layer. Each value was classified as a salient event or not using the same exponentially decaying criterion as before (see Table S3 for criterion values). Finally, accumulated salient events were mapped onto units of seconds using multiple linear regression.

## Acknowledgements

Thank you to Charlotte Rae, Petar Raykov, Samira Bouyagoub, Chris Bird, Francesca Simonelli, and Mara Cercignani for their assistance with this project. Thanks also to Virginie van Wassenhove and Martin Wiener for comments on an earlier version of the manuscript.

---

**Figure S1.**
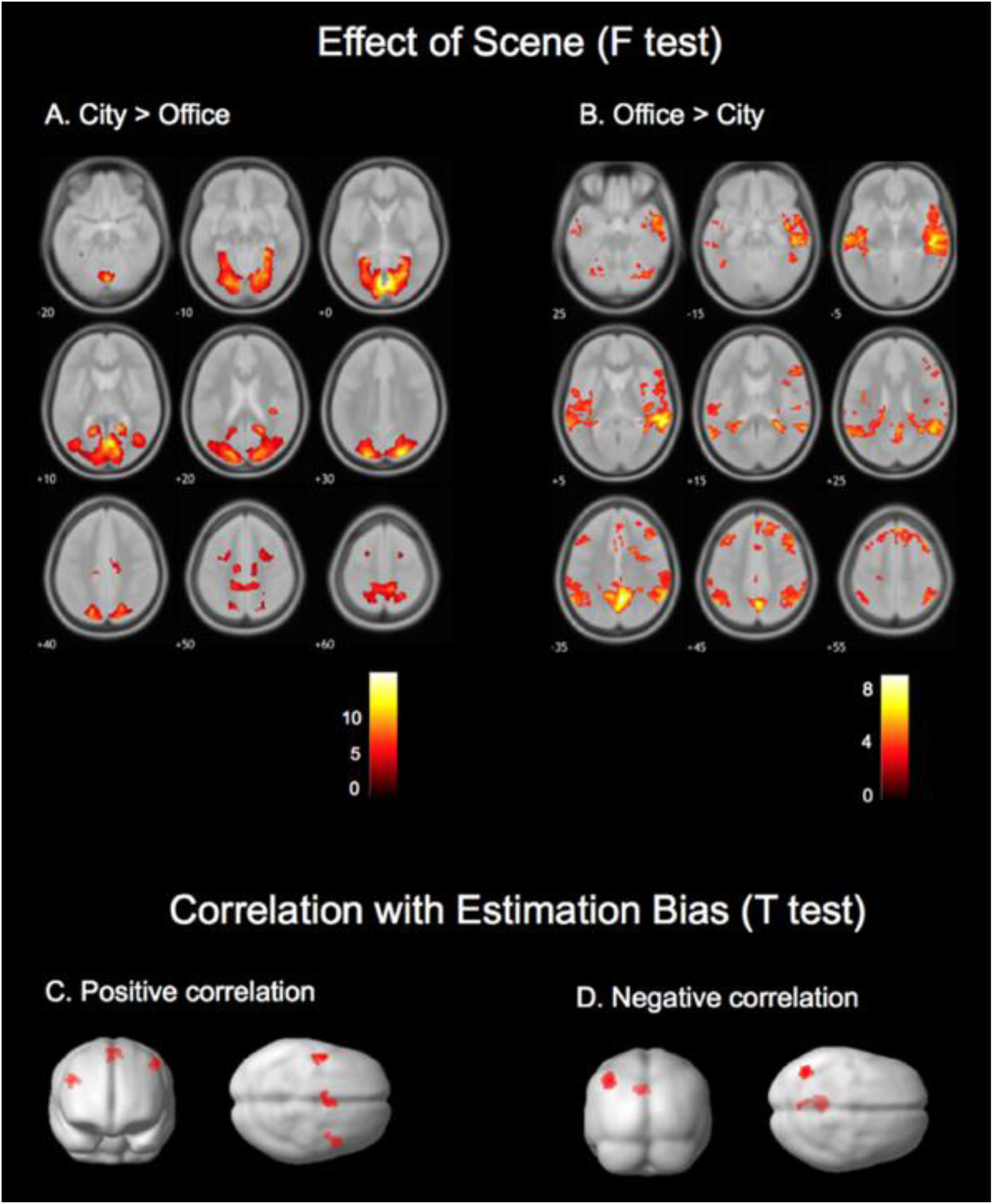
Results from confirmatory GLM on BOLD (significant clusters only). **A** Higher BOLD for city than office scenes: R lingual gyrus; bilateral midcingulate area; R insula; bilateral SFG. **B** Higher BOLD for office than city scenes: R precuneus; bilateral precentral gyrus; L MFG; bilateral cerebellum; L paracentral lobule; R SFG. **C** Positive correlation with normalized estimation bias: bilateral precentral gyrus; L SMA; R superior occipital gyrus. **D** Negative correlation with normalized estimation bias: L angular frontal gyrus; L MFG; L posterior cingulate. See also Table S4.

**Figure S2.**
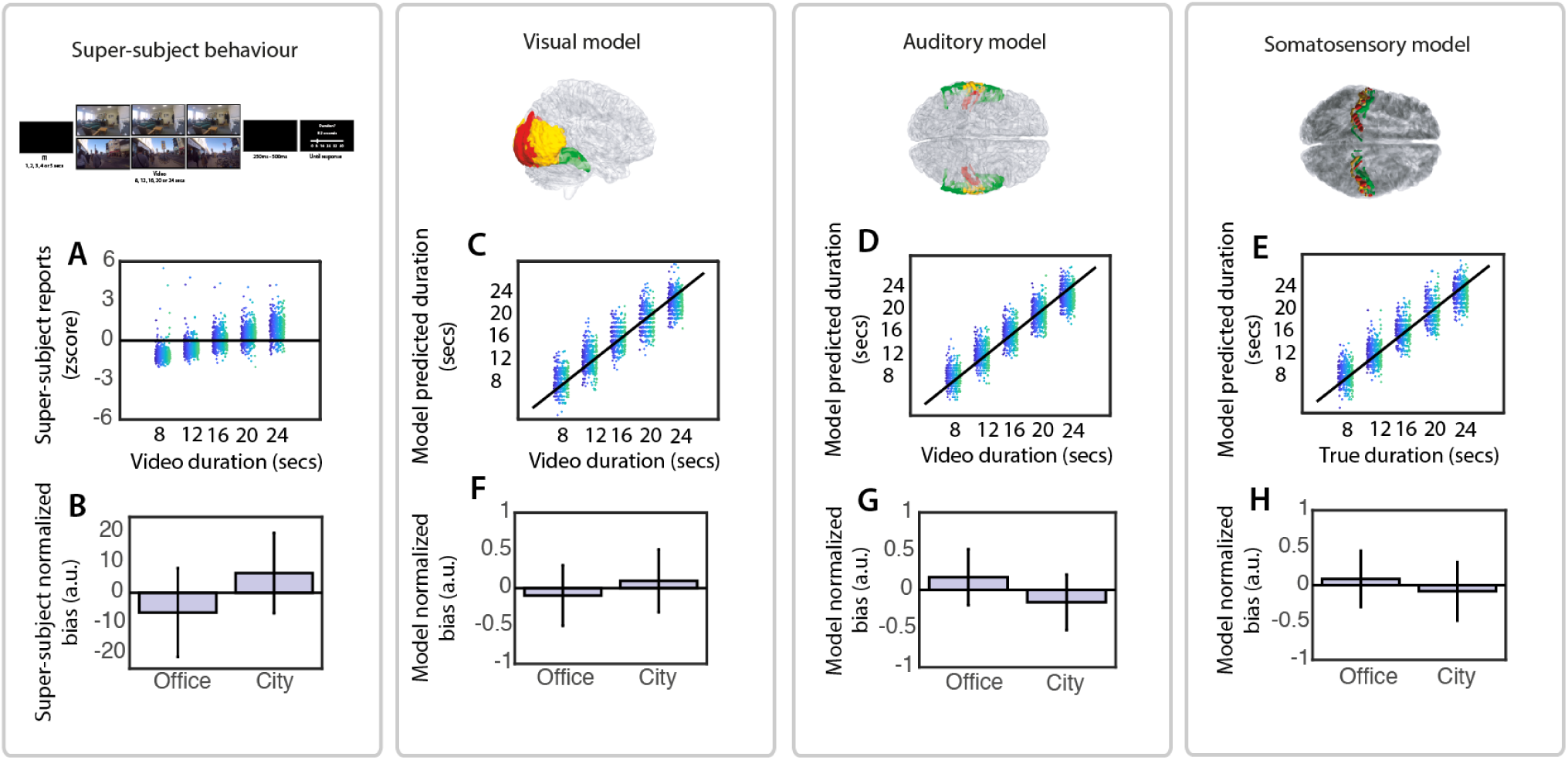
Brain-based modelling on the pre-registered pipeline. **(A)** Strong positive association between veridical video durations and the z-scored reports we used to build the super-subject. **(B)** Normalized estimation bias computed on pooled (‘super-subject’) behavioral data, as a function of video scene. **(C-E)** Association between veridical video duration and model-predicted durations separately for visual, auditory and somatosensory Euclidean Distance models respectively. **(F-H)** Mean normalized bias of the visual, auditory and somatosensory models respectively, for office versus city scenes. Dot colors in the scatterplots represent different participants. Error bars in the bar charts represent SEM.

**Figure S3.**
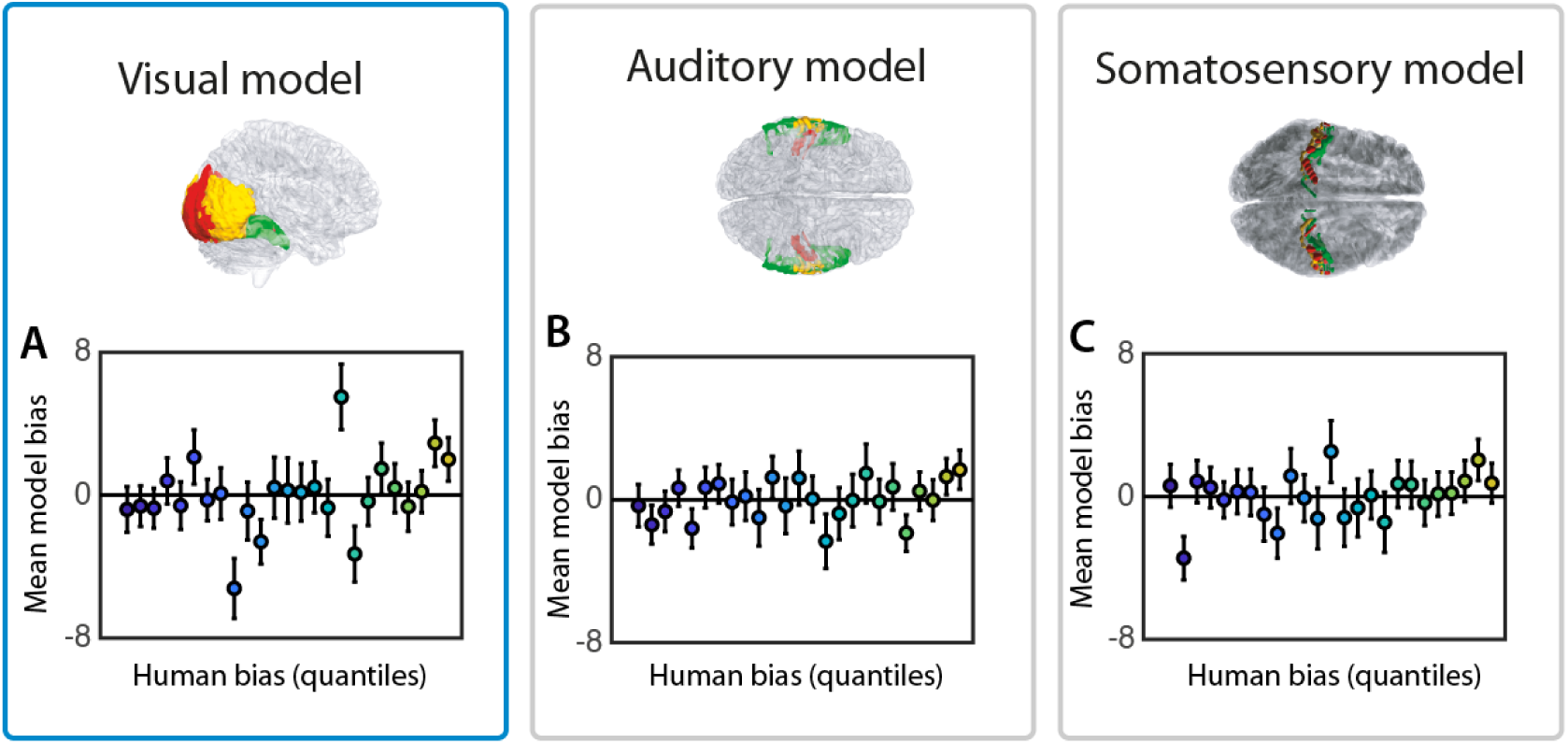
Normalized bias predicted by models trained on salient events (Euclidean distance) in **(A)** visual, **(B)** auditory and **(C)** somatosensory hierarchies. On the x-axis is the 25 bins representing 25 quantiles of human super-subject bias, and on the y-axis is mean model bias for the trials that fell within in the respective bins. Error bars represent +/- SEM.

**Figure S4.**
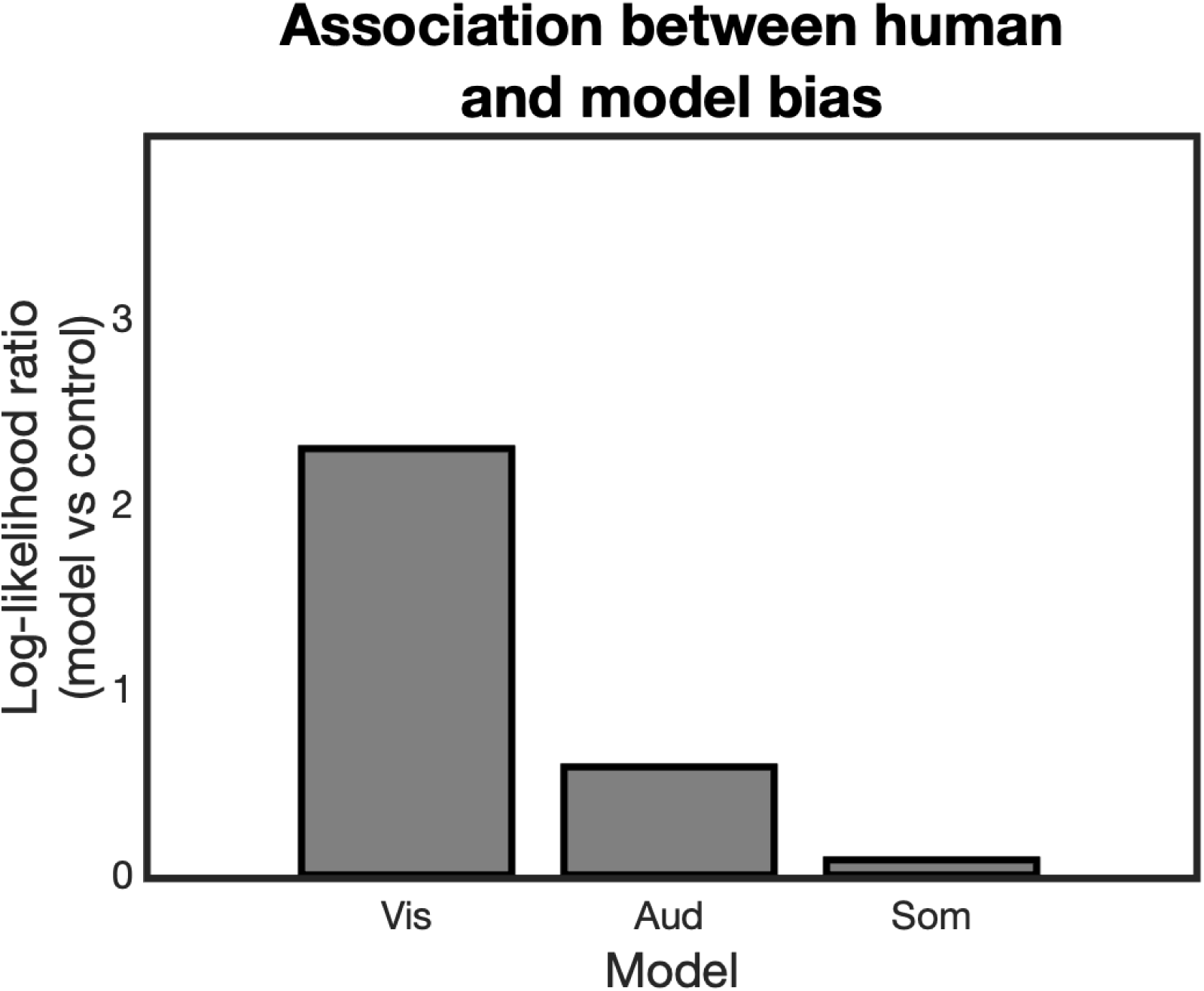
Model fits for the regression of predicted durations onto veridical durations (Euclidean Distance models), expressed as log-likelihood ratios. For each of the visual, auditory and somatosensory models, we regressed the model-predicted biases onto the human super-subject’s biases. To compare the performance of the three regressions we compared their log-likelihoods to the null (intercept) model (higher values indicate better model fits). The visual cortex regression outperforms the other two, as indicated by its higher log-likelihood ratio.

**Figure S5.**
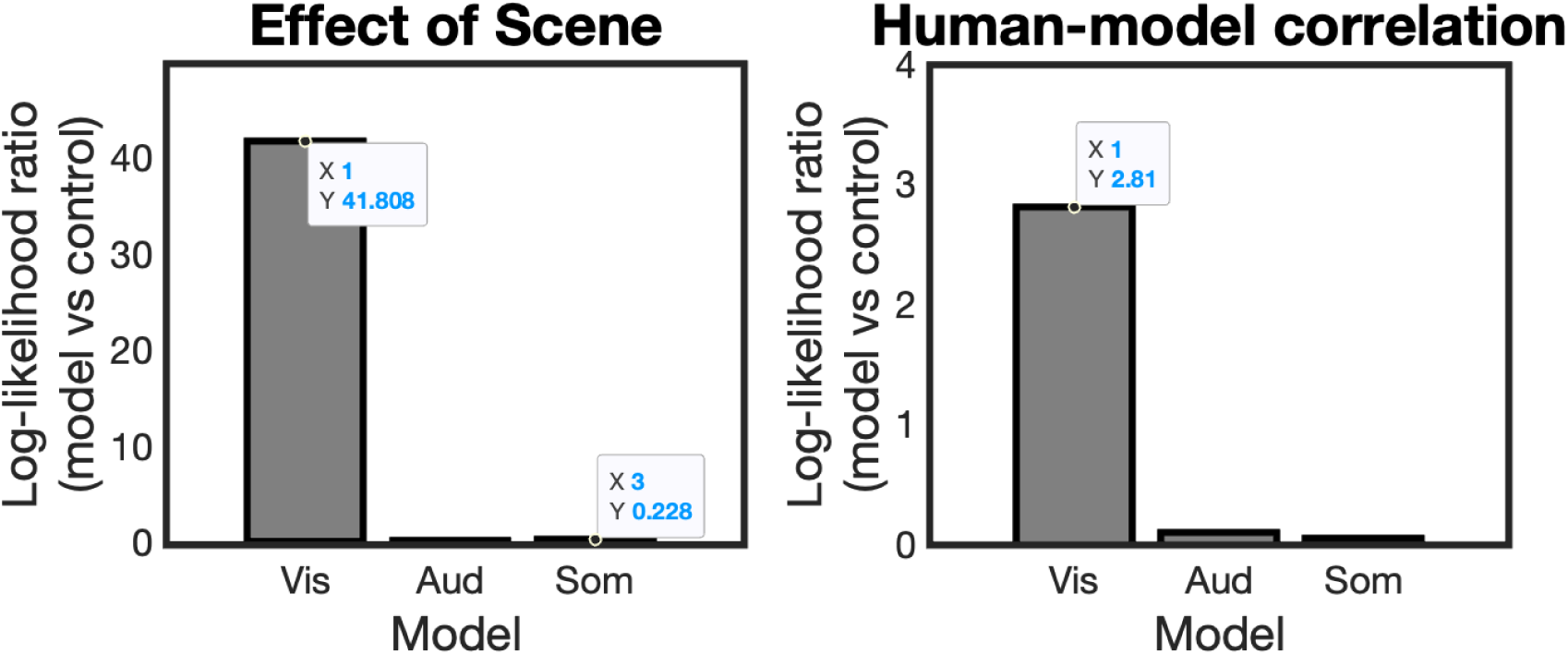
Model fits for the linear mixed models (Signed Difference analyses). **Left.** To test whether the visual, auditory or somatosensory models generated predicted durations that discriminated video type, we ran linear mixed models (LMMs) predicting model biases from the fixed effect video scene (city vs office). These were compared to control LMMs that did not have this fixed effect, using the log-likelihood ratio (LLR). The visual cortex LMM outperformed the auditory and somatosensory cortex LMMs as indicated by the greater LLR. **Right.** For each of the visual, auditory and somatosensory models, we constructed an LMM with human bias as the outcome and the model-predicted biases as a fixed effect. These LMMs tested the video-by-video correlations between predicted and human bias. These LMMs were compared to control models that did not have the model-predicted bias as a fixed effect, using LLR. The visual cortex LMM outperformed the auditory and somatosensory cortex LMMs, as indicated by the greater LLR.

**Table S1.** Definition of hierarchies for each sensory cortex model

|  | Visual | Auditory | Somatosensory |
| --- | --- | --- | --- |
| Layer<br>1 | V1, V2v, V3v | BA41 | BA3 |
| Layer<br>2 | hV4, LO1, LO2 | BA42 | BA1 |
| Layer<br>3 | VO1, VO2, PHC1, PHC2 | BA22 | BA2 |

**Table S2.** Criterion parameters for each hierarchical layer of the sensory cortex models

| Layer | $a$ | $\vartheta_{max}$ | $\vartheta_{min}$ |
| --- | --- | --- | --- |
|  |  | (SD above the mean) | (SD below the mean) |
| 1 | 0.5 | 0.5 | 1 |
| 2 | 1 | 1 | 0.5 |
| 3 | 1.5 | 1.5 | 0 |

**Table S3.** Criterion parameters for the artificial network model

| Layer | $t_{\max}$ | $t_{\min}$ |
| --- | --- | --- |
| conv1 | 39973 | 0 |
| conv2 | 11601 | 0 |
| conv3 | 5515 | 0 |
| conv4 | 3244 | 0 |
| conv5 | 1117 | 0 |
| fc6 | 124 | 0 |
| fc7 | 31 | 0 |
| output | 0.33 | 0 |
| For all layers, $\alpha = 0.001/t_{\max}$, $\tau = 6.6T_{\max}$ while training and $\tau = 10T_{\max}$ while testing. | | |

**Table S4.** Significant clusters revealed by confirmatory GLM on BOLD

| Region | Size (cm<sup>3</sup>) | *T* or *F* | *P*<sub>FWE</sub> | Peak MNI |  |  |
| --- | --- | --- | --- | --- | --- | --- |
|  |  |  |  | x | y | z |
| City > Office (two-tailed) |  |  |  |  |  |  |
| R Lingual Gyrus | 121.37 | 14.69 | < 0.001 | 6 | -70 | 4 |
| L Midcingulate Area | 1.20 | 8.86 | 0.002 | -10 | -20 | 46 |
| R Insula | 0.6 | 7.91 | 0.049 | 36 | -28 | 22 |
| R Midcingulate Area | 1.66 | 6.73 | <0.001 | 12 | -12 | 44 |
| R Superior Frontal Gyrus | 2.11 | 6.44 | < 0.001 | 24 | 0 | 54 |
| L Superior Frontal Gyrus | 1.26 | 5.94 | 0.001 | -22 | 0 | 54 |
| Office > City (two-tailed) |  |  |  |  |  |  |
| R Precuneus | 101.80 | 9.48 | < 0.001 | 6 | -56 | 36 |
| R Precentral Gyrus | 1.86 | 6.45 | < 0.001 | 24 | -26 | 66 |
| L Middle Frontal Gyrus | 4.02 | 6.42 | < 0.001 | -32 | 30 | 48 |
| R Cerebellum 1 | 2.39 | 6.12 | < 0.001 | 46 | -62 | -26 |
| L Precentral Gyrus | 1.18 | 5.82 | 0.002 | -22 | -28 | 62 |
| L Cerebellum 6 | 1.06 | 5.51 | 0.003 | -22 | -70 | -22 |
| L Paracentral Lobule | 0.82 | 4.92 | 0.013 | -6 | -12 | 68 |
| L Superior Frontal Sulcus | 0.93 | 4.44 | 0.007 | -14 | 30 | 54 |
| Positive correlation with normalized bias |  |  |  |  |  |  |
| R Precentral gyrus | 0.90 | 4.88 | 0.002 | 38 | 2 | 30 |
| L Precentral gyrus | 0.71 | 4.86 | 0.006 | -48 | 0 | 54 |
| L Supplementary motor area | 0.82 | 4.53 | 0.003 | 0 | 0 | 64 |
| R Superior Occipital Gyrus | 0.48 | 4.03 | 0.041 | 24 | -64 | 46 |
| Negative correlation with normalized bias |  |  |  |  |  |  |
| L Angular Gyrus | 1.46 | 5.54 | < 0.001 | -40 | -64 | 44 |
| L Middle Frontal Gyrus | 1.66 | 5.00 | < 0.001 | -2 | -44 | 30 |
| L Posterior Cingulate | 0.62 | 4.96 | 0.013 | -28 | 24 | 56 |

## Notes

**Conflict of interest statement:** The authors declare no conflicts of interest.

**Funding:** This work was supported by the European Union Future and Emerging Technologies grant (GA:641100) TIMESTORM – Mind and Time: Investigation of the Temporal Traits of Human-Machine Convergence and the Dr Mortimer and Theresa Sackler Foundation (MTS and AKS), which supports the Sackler Centre for Consciousness Science. AKS is also grateful to the Canadian Institute for Advanced Research (CIFAR) Program in Brain, Mind, and Consciousness.

https://osf.io/2zqfu

### Competing Interest Statement

The authors have declared no competing interest.

### Summary of Updates

- Substantial rewriting of manuscript - New analyses (see Fig. 5 and Fig. 8) - New title

